# Extra-hypothalamic oxytocin neurons drive stress-induced social vigilance and avoidance

**DOI:** 10.1101/2020.06.02.129981

**Authors:** Natalia Duque-Wilckens, Lisette Y. Torres, Sae Yokoyama, Vanessa A. Minie, Amy M. Tran, Stela P. Petkova, Rebecca Hao, Stephanie Ramos-Maciel, Roberto A. Rios, Kenneth Jackson, Francisco J. Flores-Ramires, Israel Garcia-Carachure, Patricia A. Pesavento, Sergio D. Iñiguez, Valery Grinevich, Brian C. Trainor

## Abstract

Oxytocin increases the salience of both positive and negative social contexts and it is thought that these diverse actions on behavior are mediated in part through circuit-specific action. This hypothesis is based primarily on manipulations of oxytocin receptor function, leaving open the question of whether different populations of oxytocin neurons mediate different effects on behavior. Here we inhibited oxytocin synthesis in a social stress-sensitive population of oxytocin neurons specifically within the medioventral bed nucleus of the stria terminalis (BNSTmv). Oxytocin knock-down prevented stress-induced increases in social vigilance and decreases in social approach. Viral tracing of BNSTmv oxytocin neurons revealed fibers in regions controlling defensive behaviors including lateral hypothalamus, anterior hypothalamus, and anteromedial BNST (BNSTam). Oxytocin infusion into BNSTam in stress naïve mice increased social vigilance and reduced social approach. These results show that a population of extra-hypothalamic oxytocin neurons play a key role in controlling stress-induced social anxiety behaviors.

## Introduction

Social anxiety disorder (SAD) is one of the most prevalent and debilitating psychiatric disorders across nations (Beesdo-Baum et al., 2012; Kessler et al., 2005; Ruscio et al., 2008; Stein et al., 2017) and is characterized by a persistent fear or anxiety of unfamiliar social situations or scrutiny by others (American Psychiatric Organization, 2013). Available treatments for SAD are ineffective for about half of the patients (Ipser et al., 2008; Liebowitz et al., 2005; Van Ameringen et al., 2001), which highlights the need for a better understanding of the underlying neurobiological alterations. A key region mediating fear and anxiety is the bed nucleus of the stria terminalis (BNST), a highly complex component of the extended amygdala that has increasingly received attention as an important contributor to stress-induced psychiatric disorders (Avery et al., 2016; Goode and Maren, 2017; Lebow and Chen, 2016). Recent evidence shows that compared to healthy controls, individuals diagnosed with SAD show increased phasic activation of the BNST in anticipation of aversive events (Figel et al., 2019), and altered BNST functional connectivity following unpredictable threats (Clauss et al., 2019). Nonetheless, it is unclear at a mechanistic level how the BNST mediates behavioral phenotypes related to SAD.

One central phenotype contributing to SAD etiology and maintenance is biased attention to social threat, which can be expressed by (1) attentional *avoidance* of socially salient information and/or (2) exacerbated *vigilance* of the social environment (Chen and Clarke, 2017; Heimberg, 1995; Spence and Rapee, 2016). Remarkably, social stress can induce both behavioral analogs in rodents (Duque-Wilckens et al., 2018; Newman et al., 2019), providing a unique opportunity to assess the neurobiological substrates underlying behavioral phenotypes associated with SAD. Using the California mouse (*Peromyscus californicus*), we recently found that stress-induced reductions in social approach and increases in social vigilance are mediated by the BNST. Oxytocin is an important regulator of social behaviors (Bosch and Young, 2018; Grinevich et al., 2015; Rogers-Carter et al., 2018; Veenema and Neumann, 2008), and it has been hypothesized that oxytocin increases the salience of both positive and negative social stimuli (Shamay-Tsoory and Abu-Akel, 2016) through distinct neural circuits (Steinman et al., 2019). In female California mice, social defeat induces hyperactivity of oxytocin neurons in medioventral BNST (BNSTmv) up to 10 weeks after the last episode of social defeat (Steinman et al., 2016). Pharmacological blockade of oxytocin receptors in the anteromedial BNST (BNSTam), but not the ventral striatum, prevents stress-induced decreases in social approach and increases vigilance (Duque-Wilckens et al., 2018).

An outstanding question is whether oxytocin produced within the BNST is necessary for stress-induced social deficits. Here we use oligonucleotide analogs to inhibit oxytocin synthesis, viral tracing to identify axonal projections of BNSTmv oxytocin neurons, and site-specific infusion of oxytocin to show that a novel oxytocinergic circuit within BNST is a key mediator of social avoidance and vigilance. Thus, while oxytocin acting in hypothalamic-mesolimbic circuits promotes social approach (Dölen et al., 2013; Marlin et al., 2015; Song et al., 2016; Wang and Aragona, 2004; Williams et al., 2020), our data in the BNST support the hypothesis that divergent effects of oxytocin on social approach are mediated by discrete brain circuits (Grinevich et al., 2016; Shamay-Tsoory and Abu-Akel, 2016; Steinman et al., 2019).

## Results

### Oxytocin antisense in BNSTmv prevents stress-induced reduction in social approach and increased vigilance

In previous studies, social defeat increased oxytocin mRNA and oxytocin/c-fos colocalizations in cells of the BNSTmv in females but not males following a social interaction test (Steinman et al., 2016). These phenotypes are accompanied by reduced social approach and increased vigilance, and are independent of estrous stage (Duque-Wilckens et al., 2018; Greenberg et al., 2013; Trainor et al., 2013; Williams et al., 2018). To test whether stress-induced increases in oxytocin synthesis within the BNSTmv are necessary for stress-induced social deficits, females were first randomly assigned to social defeat or control conditions (Fig. 1B). One week later we made a single infusion of vivo morpholinos targeting Oxt mRNA (antisense) or missense into the BNSTmv (Fig.1A). Overall, antisense reduced the number of oxytocin-immunoreactive cells detected in the BNSTmv compared to missense controls (F_1,28_=5.4, p=0.02, Fig.1C, 1D). Planned comparisons revealed that compared to missense, antisense reduced oxytocin immunoreactive cells in stressed (p=0.03, d=1), but not in control females. There was no evidence of neurotoxicity by morpholino treatment (supplementary fig. 1B), consistent with previous reports (Reissner et al., 2012). There were no effects of BNST antisense infusions on PVN oxytocin cells (supplementary fig. 1F), indicating oxytocin knock-down in the BNST was site-specific. An alternate schedule of multiple injections of a lower dose of morpholino antisense also reduced oxytocin cell number in the BNSTmv (stress missense vs. stress antisense p=0.01, d=1.16, supplementary fig. 1H,1I).

**Fig. 1.**
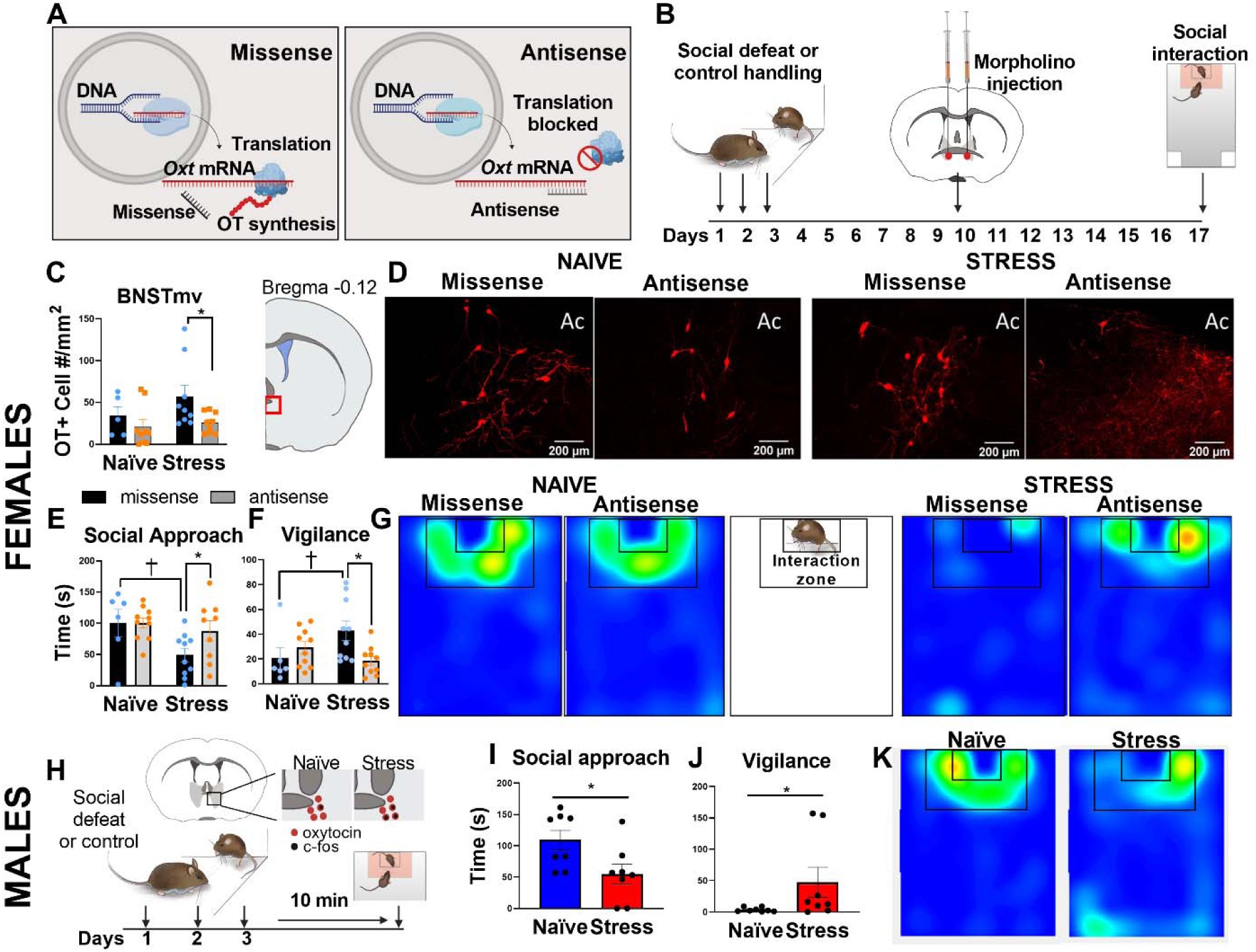
Oxytocin synthesis within BNSTmv is necessary for stress-induced increases in social vigilance and reduced social approach. **A**. Morpholino oligos inhibit protein synthesis by blocking the translation initiation complex. **B**. Timeline of experiment assessing the effects of oxytocin antisense injections in BNSTmv on oxytocin cell number and behavior. **C**. Morpholino antisense reduced the number of oxytocin+ cells detected in BNSTmv compared to missense controls (n=5-10 per group). **D**. Representative images of BNSTmv oxytocin (+) cells in naïve and stressed animals receiving missense vs. antisense injections. **E**,**F**. Effects of social defeat on behavior in the social interaction test (n=6-10 per group). Social defeat reduced social approach and increased social vigilance in missense treated females but not antisense treated females. **G**. Representative heatmaps for the interaction phase showing reduced time spent in the interaction zone in females receiving missense but not antisense. **H**. Timeline of experiment assessing the behavior of male California mice 10 min after a third episode of social defeat. We previously found that at this moment, males show increased cFos/oxytocin colocalizations in the BNSTmv (Steinman et al., 2016). **I**. Acute effects of a third episode of social defeat on male behavior in the social interaction test (n=8 per group). Defeat reduced social approach and increased vigilance (**J**). **K**. Representative heatmaps for the interaction phase showing reduced time spent in the interaction zone in stressed compared to naïve males. * p<0.05 antisense vs. missense, † p<0.05 control vs. stress. Ac=anterior commissure

Mice were tested in a social interaction test with a same-sex unfamiliar target mouse. The effect of social defeat on social approach, measured by time spent in the interaction zone, was dependent on antisense treatment (F_1,31_=5.7, p=0.02, fig.1E). Defeat reduced social approach in females receiving missense (p=0.02, d=1.1), but not in females receiving antisense. For social vigilance, effects of stress were blunted by antisense treatment (stress*treatment F_1,31_=6.2, p=0.02, fig. 1F). Social vigilance was defined as time spent oriented towards the target mouse while outside of the interaction zone (Duque-Wilckens et al., 2018; Newman et al., 2019). Stress increased social vigilance in missense treated females (p=0.03, d=0.96) but not antisense treated females. There were no effects of stress or treatment on approach behavior during the acclimation phase when the target mouse was absent (supplementary fig. 1C), or time spent in center (supplementary fig. 1D) and distance traveled (supplementary fig. 1D) during the open field phase. Together, these data show that stress-induced synthesis of oxytocin in the BNST is necessary for stress-induced social deficits in female California mice.

### Male California mice show social avoidance and vigilance immediately after social defeat

Although the BNST is sexually dimorphic (Allen and Gorski, 1990; Campi et al., 2013) and known to mediate sex differences in behavior (Duque-Wilckens et al., 2016; Janitzky et al., 2014; Whylings et al., 2020), the population of oxytocin neurons within the BNSTmv is conserved in both sexes (Steinman et al., 2016). While male California mice do not show social deficits or increased oxytocin/c-fos colocalizations *two weeks* after social defeat stress (Duque-Wilckens et al., 2018; Greenberg et al., 2013; Trainor et al., 2013), males show increased oxytocin/cFos colocalizations in the BNST *immediately after* a third episode of defeat (Steinman et al., 2016). Thus, we hypothesized that social defeat stress would induce an acute decrease in social approach and increase in vigilance in males. When stressed males were tested 10 min after a third episode of defeat (Fig.1H), social approach was reduced (t_14_=2.46, p=0.02, d=1.23, fig. 1I), and social vigilance was increased (t_14_=2.36, p=0.03, d=1.18, fig.1J) compared to naïve males. There were no effects of defeat on approach behavior when the target mouse was absent (supplementary fig. 2B) or in the open field (supplementary fig. 2C, 2D). These results implicate oxytocin in the BNST as a potential modulator of social approach in males as well as females. Next, we assessed whether stress effects on the BNST extend to an alternate model species.

### In *Mus musculus*, vicarious social stress increases social vigilance and BNST OT cell number in females

Low aggression in female C57BL6/J mice prevents the use of conventional social defeat protocols (but see (Takahashi et al., 2017)), so we applied vicarious social defeat stress, which was previously shown to induce depression- and anxiety-like behaviors in both male and female mice (Iñiguez et al., 2018; Warren et al., 2020). Male and female C57BL/6J mice that experienced vicarious social defeat stress or control handling were tested in a social interaction test (Fig.2A). The effects of stress on social vigilance were stronger in females than males (fig.2B, sex*stress interaction, F_1,36_=8.2, p=0.01). Planned comparisons revealed that vicarious defeat stress significantly increased vigilance in female (p<0.0001, d=1.9), but not male mice (p=0.1, d=1.4). Immunohistochemical analyses of oxytocin cells in females showed that the distribution of oxytocin positive cells within the BNST was limited to ventral BNST near bregma 0.02 (fig. 2C), similar to California mice (Steinman et al., 2016) and rats (DiBenedictis et al., 2017). Importantly, vicarious stress increased the number of oxytocin cells in the BNST (t_14_=2.721, p=0.01, d=1.4, fig 2D), but not in the PVN (fig. 2E), comparable to what was previously reported in female California mice. Together, these data suggest that the impact of social stress on BNST oxytocin neurons is conserved across rodent species and prompted us to further investigate the circuit.

**Fig. 2.**
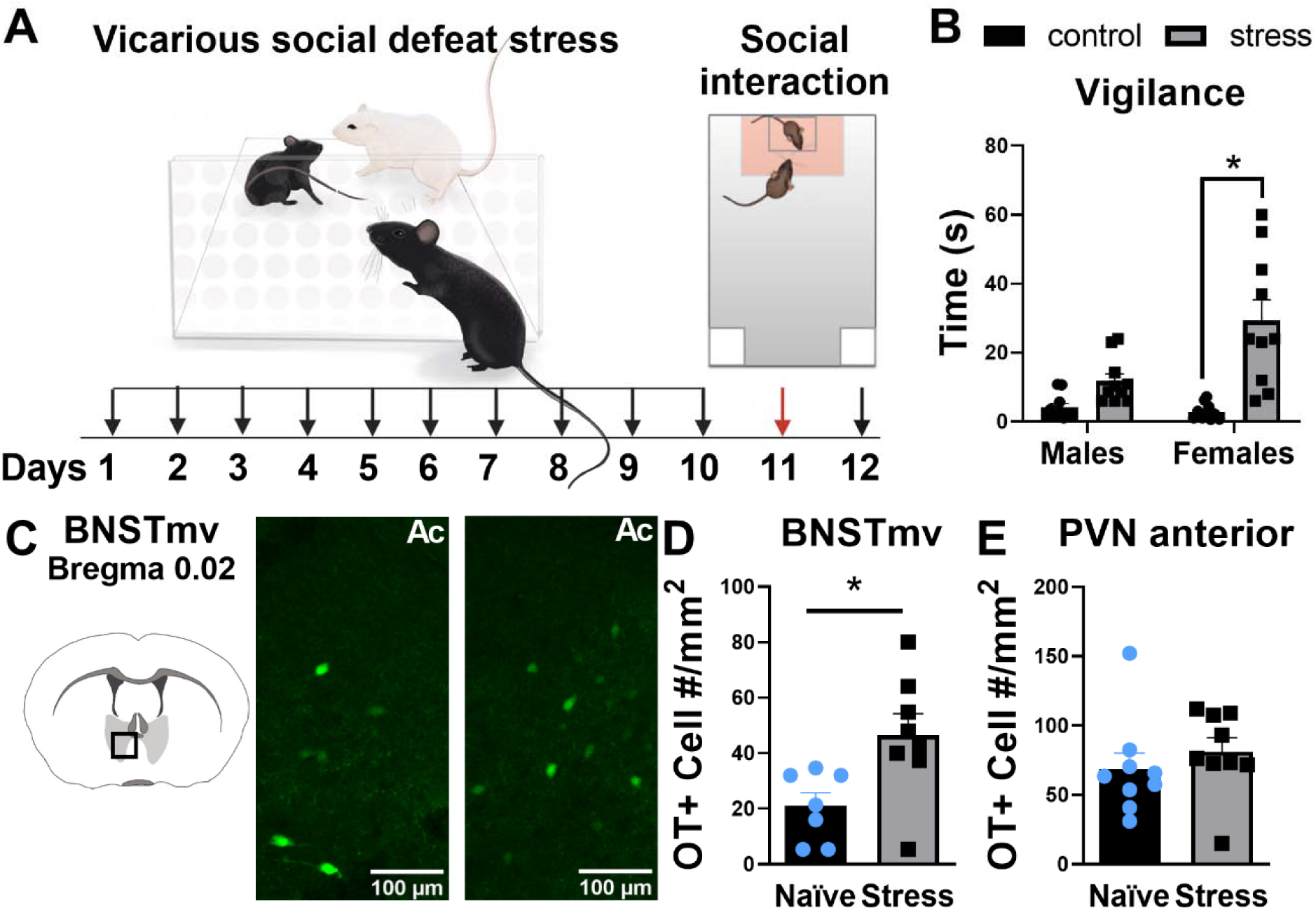
In *Mus musculus*, vicarious social stress increases social vigilance and BNST OT cell number in females. **A**. Timeline of experiment assessing effects of vicarious defeat stress on behavior and BNSTmv oxytocin cell number in *Mus musculus*. **B**. Effects of vicarious social defeat on vigilance behavior (n=10 per group). Defeat significantly increases vigilance behavior in females, but not males. **C**. Representative images of the effects of vicarious social defeat on oxytocin cell number in the BNSTmv. **D**,**E**. Effects of vicarious defeat stress on oxytocin cell number (n=7 per group). Stress significantly increased oxytocin cell number in BNST, but not PVN. * p<0.05

### Oxytocin cells in the BNSTmv project to the anteromedial BNST

To identify the anatomical projections of the oxytocin neuron population in the BNSTmv, we used an AAV construct expressing Venus under the control of mouse oxytocin promoter (Knobloch et al., 2012) in males and females (fig.3A,B). Immunohistochemical analyses in the BNST (fig. 3C) and the PVN (supplementary fig. 3A) showed that on average, 63% of oxytocin-immunoreactive cells expressed Venus and that all Venus positive cells co-expressed oxytocin. When Venus expression was exclusively limited to BNSTmv neurons, we found that in both males and females BNSTmv oxytocin neurons sent fibers to more anterior areas of the BNST including dorsal (fig. 3D) and ventral (fig. 3E) subregions of the BNSTam. We also observed fibers in hypothalamic areas including anterior fig. 3F) and lateral (Fig. 3G) hypothalamus. The identification of fibers in the dorsal BNSTam was notable, as we previously showed that activation of oxytocin receptors in dorsal BNSTam is necessary for stress-induced social deficits in females (Duque-Wilckens et al., 2018). Since BNST oxytocin neurons project to this area in both sexes, we then asked the question whether differences in expression of oxytocin receptors in the BNSTam could be a contributor to sex-specific effects of social defeat stress on behavior.

**Fig. 3.**
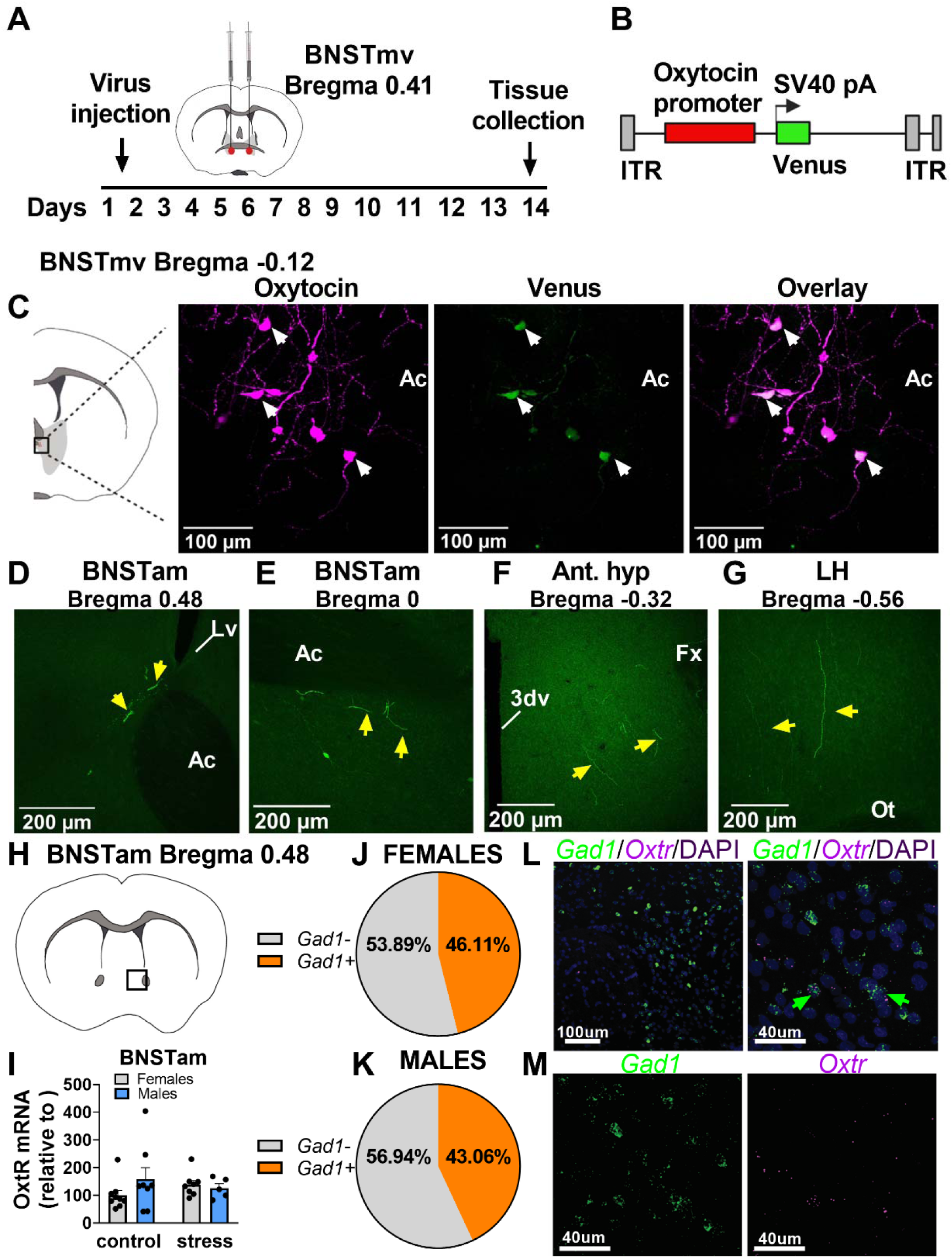
Oxytocin cell bodies in BNSTmv send projections to anteromedial BNST. **A**. Timeline of experiment using viral tracing to identify axonal projections of BNSTmv oxytocin neurons. **B**. AAV construct. **C**. Representative images showing colocalization of oxytocin and Venus in the BNSTmv. **D**,**E**,**F**,**G**. In animals which Venus expression was exclusively limited to BNSTmv neurons, Venus+ fibers were detected in hypothalamic nuclei and anterior BNST, including BNSTam. **H**. Diagram showing localization of samples taken for gene expression analyses. **I**. OxtR expression in BNSTam after social defeat (n=5-8 per group). No effects of stress or sex were found. **J**,**K**. Circle chart representing percentage of *Oxt* +/*Gad*+ cells present in the BNSTam of males and females. **L**,**M**. Representative photomicrographs of *Gad1/Oxtr*/DAPI fluorescent in situ hybridization in BNSTam. Ac=anterior commissure; Lv=lateral ventricle; 3dv=third ventricle; Fx=fornix; Ot=optic tract

### About half of BNSTam *Oxtr* neurons coexpress inhibitory markers

First, we used real-time PCR to assess whether there were basal or stress-induced sex differences in *Oxtr* expression within the BNSTam and there were no differences (fig., 3I). Next, we employed the *in situ* hybridization using probes against *Oxtr, Gad1*, and *vGlut2* to assess cell types of *Oxtr+* in the BNSTam. About half (46% females, 43% males) of the *Oxtr+* cells in the BNSTam co-expressed *Gad1+* (fig. 3J,K). We found no *vGlut2+* cells in the BNSTam, although there were *vGlut2+* cells in the adjacent lateral septum (supplementary fig. 3B). Next, we tested whether administration of oxytocin within the BNSTam is sufficient to reduce social approach and increase vigilance in stress naïve females and males (fig. 4A).

**Fig. 4.**
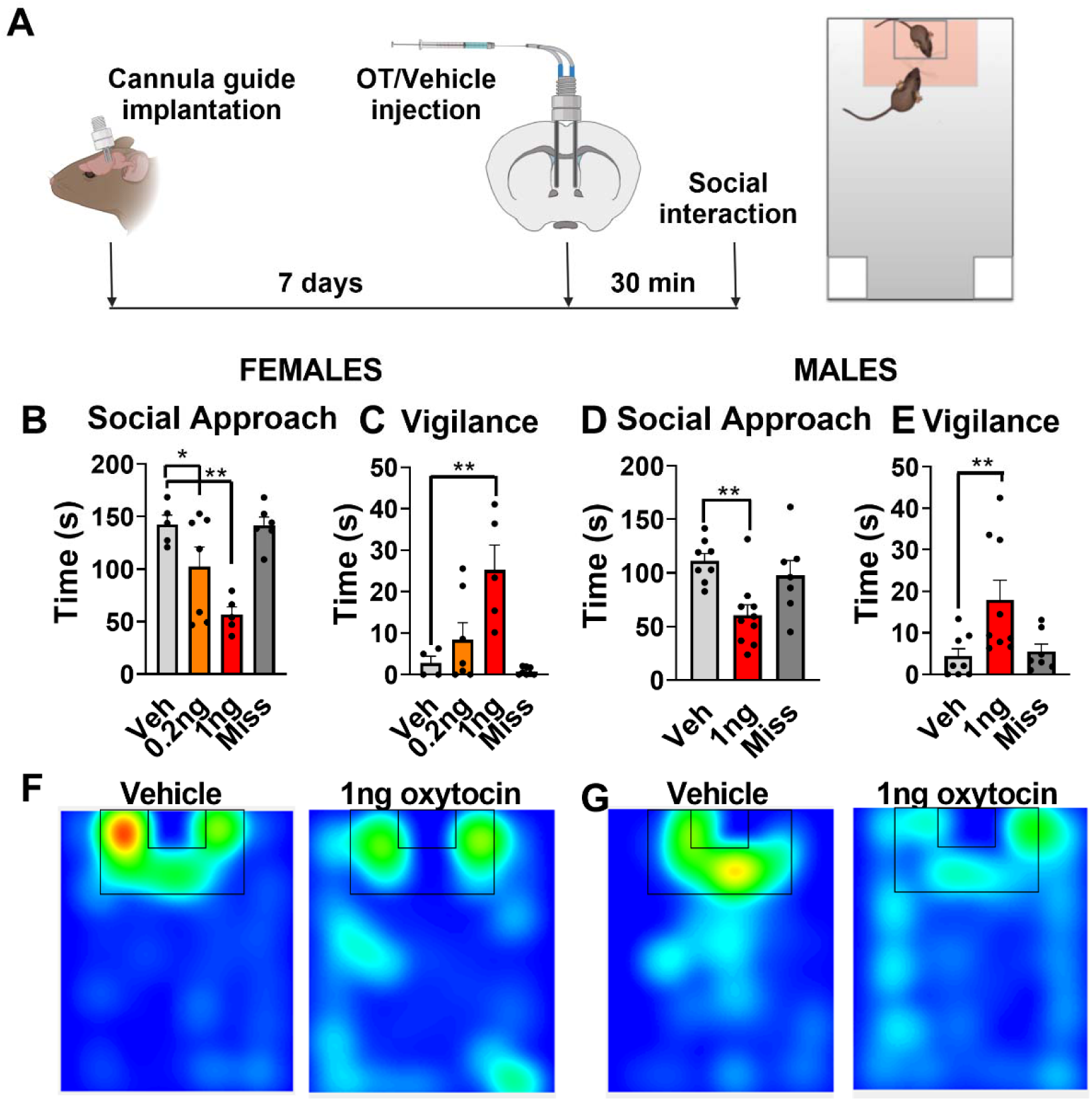
Oxytocin infusion within BNSTam induces social vigilance in stress naïve females and males. **A**. Timeline of experiment assessing behavioral effects of oxytocin injection in BNSTam in animals naïve to defeat. **B**,**C**. Effects of 0.2ng or 1ng injections of oxytocin within BNSTam on female behavior in the social interaction test (n=5-7 per group). Oxytocin dose dependently reduced social approach, and only the high dose of oxytocin increased vigilance. **D**,**E**. Effects of 1ng injection of oxytocin within BNSTam on male behavior in the social interaction test (n=6-9 per group). Oxytocin reduced social approach and increased vigilance in males. **F**,**G**. Representative heatmaps for the interaction phase showing reduced time spent in the interaction zone in naïve males and females receiving oxytocin injections in BNSTam. * p<0.05, **p<0.01

### Oxytocin infusion within the BNSTam induces social vigilance in stress naïve female and male mice

In females, treatment with oxytocin decreased social approach (F_3,21_=9.5, p<0.001, fig. 4B). Compared to vehicle, both 0.2 ng (p=0.03, d=1.1) and 1.0 ng (p<0.001, d=4.8) infusions of oxytocin reduced social approach in a dose-dependent fashion. Treatment with oxytocin also increased social vigilance in females (F_3,21_=8.8, p=0.001, fig. 4C). The 1.0 ng (p<0.001, d=2.3) but not the 0.2 ng dose increased vigilance in stress naïve females compared to vehicle. There were no effects of oxytocin injections placed outside the BNSTam (miss) on social approach or vigilance. In males, oxytocin also reduced social approach (F_2,21_=6.7, p=0.00), with 1.0 ng of oxytocin reducing social approach compared to vehicle (p=0.00, d=1.93 fig. 4D). Similarly, oxytocin also increased social vigilance (F_2,21_=5.3, p=0.01, fig. 4E), with 1.0 ng of oxytocin increasing vigilance compared to vehicle (p<0.001, d=1.27). There were no effects of oxytocin injections placed outside the BNSTam on social approach or vigilance. Together, our results suggest that the oxytocin-dependent circuit of social vigilance in the BNST is intact in both sexes, and that previously reported sex differences in behavioral responses to social stress (Duque-Wilckens et al., 2018; Trainor et al., 2011) are mediated by sex differences in the release of oxytocin.

## Discussion

The social salience hypothesis proposed that oxytocin increases attention and motivational responses to social contexts (Bartz et al., 2011; Shamay-Tsoory and Abu-Akel, 2016). However, the underlying mechanisms that allow oxytocin to have both anxiolytic (Blume et al., 2008; Domes et al., 2007; Kirsch et al., 2005; Knobloch et al., 2012) and anxiogenic (Eckstein et al., 2014; Martinon et al., 2019; Nasanbuyan et al., 2018) effects have remained largely elusive. Here we outlined an oxytocinergic circuit within the BNST that is a critical hub driving stress-induced changes in social behavior. Antisense knockdown of oxytocin synthesis within the BNSTmv showed that this group of neurons is necessary for long-term decreases in social approach and increased vigilance resulting from exposure to social defeat. Viral tracing showed that BNSTmv oxytocin neurons project to the BNSTam, a region in which oxytocin receptor inhibition is sufficient to prevent stress-induced social deficits (Duque-Wilckens et al., 2018).. Finally, oxytocin infusion into the BNSTam was sufficient to reduce social approach and increase social vigilance in males and females naïve to defeat. Together, these data suggest that stress-sensitive oxytocin neurons located in the BNSTmv project to the BNSTam, where oxytocin acts to drive a social anxiety-related phenotype. To our knowledge, this is the first demonstration that non-hypothalamic oxytocin synthesis can modulate behavior.

### Oxytocin produced by BNST neurons promotes social vigilance and reduces social approach

The BNST is a known modulator of anxiety-related behaviors (Fox and Shackman, 2019; Lebow and Chen, 2016), and recent imaging data in humans showed that BNST activity is altered in patients suffering from SAD (Clauss et al., 2019; Figel et al., 2019). Previous work manipulating oxytocin receptor activity showed that oxytocin acting in the BNST can facilitate expression of anxiety-like behaviors (Martinon et al., 2019; Moaddab and Dabrowska, 2017), but the source of oxytocin was not clear. Using antisense knockdown, we show that oxytocin *synthesized* in neurons within the BNST is necessary for stress-induced social vigilance and reduced social approach. Although the presence of oxytocin cells in the BNST has been described in multiple species (Caffé et al., 1989; DiBenedictis et al., 2017; Nasanbuyan et al., 2018), this is the first time that a behavioral function of these neurons has been demonstrated. Social stress increases oxytocin cell mRNA and counts in the BNST of female California mice (Steinman et al., 2016) and C57BL6/J mice. In California mice, social defeat induces a delayed increase in reactivity of BNST oxytocin neurons (Steinman et al., 2016) and a reduction in social approach in females but not males (Duque-Wilckens et al., 2018; Greenberg et al., 2014; Steinman et al., 2016). An intriguing question is why the long-term effects of defeat on those BNST oxytocin neurons are stronger in females than in males.

One possible mechanism is brain derived neurotrophic factor (BDNF). Defeat increases BDNF protein within the BNST two weeks after social defeat in females but not males, and defeat-induced social avoidance is dependent on TrkB signaling in the BNST (Greenberg et al., 2014). In the hypothalamus, BDNF signaling through TrkB is necessary for oxytocin-dependent maternal behavior (Maynard et al., 2018), suggesting that BDNF-TrkB signaling may also enhance the anxiogenic effects of oxytocin in the BNST. This could occur through enhancement of long-term potentiation in oxytocin neurons, which is enhanced by BDNF through pre- and post-synaptic mechanisms (Park and Poo, 2013). Additionally, BDNF could increase synapsis formation, which occurs in hypothalamic neurons during the peripartum period resulting in increased excitability (Hatton, 1990; Stern and Armstrong, 1998; Theodosis et al., 2004). These mechanisms could account for why BNST oxytocin neurons are still more reactive in females up to ten weeks after social defeat. Epigenetic mechanisms might contribute as well. Sex-specific effects on stress on DNA methyltransferase expression have been reported in other brain regions such as the nucleus accumbens (Hodes et al., 2015) and central nucleus of the amygdala (Wright et al., 2017). Curiously, after a third episode of defeat, male California mice had elevated oxytocin/c-fos colocalization in the BNST (Steinman et al., 2016). We found that comparable to females, stressed male California mice show reduced social approach and increased social vigilance at this time point. Acute increases in BNST oxytocin/c-fos colocalization has also been described immediately after social defeat in male C57BL6/J mice (Nasanbuyan et al., 2018). This suggested that the behavioral function of the BNST oxytocinergic neural circuitry may be conserved between sexes. We considered this question when we explored the descending pathways of BNST oxytocin neurons.

### BNST oxytocin neurons project to nuclei that control defensive behavior and social vigilance

Viral tracing experiments in males and females revealed fibers originating from oxytocin neurons within the BNST, anterior hypothalamus, and lateral hypothalamus, consistent with previous tracing experiments (Delville et al., 2000; Dong and Swanson, 2004) that were not cell-type specific. Although we do not know whether BNST oxytocin neurons synapse with neurons within these regions, there is evidence for non-synaptic, periaxonal or *en passant* release of oxytocin (Chini et al., 2017; Knobloch and Grinevich, 2014). Thus, a nucleus containing only fibers of passage could receive oxytocin even in the absence of axon terminals. Interestingly, there were many fibers in the BNSTam, which is implicated as a key regulator of social anxiety-related behavior. Social anxiety phenotypes are associated with increased activity in the BNSTam (Duque-Wilckens et al., 2018; Kollack-Walker et al., 1997; Markham et al., 2009), and pharmacological inhibition of oxytocin receptors in the BNSTam restores normative social approach behavior following social defeat (Duque-Wilckens et al., 2018). Venus positive fibers in the BNSTam originating from the BNSTmv were observed in both sexes, which further supported the hypothesis that the circuitry for social vigilance is intact in males and females. Indeed, infusion of oxytocin into the BNSTam reduced social approach and increased social vigilance in both males and females naïve to social defeat. Together, these results suggest that sex differences in stress-induced vigilance are mediated by sex-specific temporal activation and plasticity of oxytocinergic BNST neurons.

About 50% of *Oxtr+* cells in the BNSTam coexpress *Gad1+*, suggesting that oxytocin may act on inhibitory neurons in the BNST as has been observed in the central nucleus of amygdala (Knobloch et al., 2012). Many *Oxtr+* cells did not express *Gad1+*. There were no *vGlut2* positive cells observed in the BNSTam, so *Oxtr+/Gad1*-cells may instead express *Gad2* (Welch et al., 2019) or represent another type of neuronal or non-neuronal cells such as glia (Mittaud et al., 2002). Future single cell analysis of the BNSTam will be highly informative in understanding how activation of *Oxtr* modulates social approach and vigilance.

### Functional implications

Previous work indicated that different circuits mediated positive (Han et al., 2018; Hung et al., 2017; Knobloch et al., 2012; Ross et al., 2009; Tang et al., 2020) and negative (Duque-Wilckens et al., 2018; Steinman et al., 2016) effects of oxytocin on social approach. A lingering question was how these circuits could be distinctly activated considering that PVN oxytocin neurons project widely throughout the brain (Knobloch and Grinevich, 2014). Our results show that stress-induced intra-BNST oxytocin synthesis facilitates social anxiety-related behaviors. Our results do not exclude a role of the PVN for inducing social vigilance, as volume diffusion of oxytocin through extracellular space or the ventricular system gives the PVN an extended reach (Landgraf and Neumann, 2004). However, the key role of BNST oxytocin neurons play in long term changes in female-biased social avoidance and vigilance suggests that these cells may be related to symptoms associated with social anxiety disorder (Asher et al., 2017) and depression (Kessler et al., 2005), which are more common in women than men. In sum, data in multiple species support an important role of the BNST in social anxiety-related behaviors, and our work shows that oxytocin signaling may be a key underlying mechanism.

## Materials and methods

### Experiments in California mice

#### Animals

All studies were approved by the Institutional Animal Care and Use Committee (IACUC) and conformed to NIH guidelines. California mice (*Peromyscus californicus*) were bred at UC Davis and group housed after weaning in clear polypropylene cages containing sanichip bedding, nestlets, and enviro-dri. Mice were housed on a 16L:8D light:dark cycle (lights off at 1400), and water and food (Harlan Teklad 2016; Madison, WI) were provided ad *libitum*. All experimental mice were at least 90 days old.

#### Social defeat stress

Mice were randomly assigned to control handling or social defeat for 3 consecutive days as described previously (Trainor et al., 2011). Control mice were placed in a clean cage for 7 min. Mice assigned to social defeat were placed in the home cage of an aggressive same-sex mouse. Each episode lasted 7 min or until the resident attacked the focal mouse 7 times (whichever occurred first). No animal was physically injured. Immediately after defeat or control handling, mice were returned to their home cage. All behavioral and brain tissue analyses were conducted two weeks after the last episode of social defeat.

#### Social Interaction test

This test consisted of three phases lasting 180 seconds each. In the open field phase, the focal mouse was introduced into an empty arena (89 × 63 x 60 cm)(Duque-Wilckens et al., 2018). In the acclimation phase, an empty wire cage was placed against one wall of the arena. In the interaction phase, an unfamiliar same-sex adult mouse was placed inside the wire cage. Total distance traveled and time spent in a) the center of the arena (within 8 cm of the sides and within a center zone located 14 cm from the sides) and b) within 8 cm of the cage (interaction zone) were recorded and analyzed via an automated video-tracking system (Anymaze, Stoelting). Vigilance behavior, defined as time spent with head oriented toward the target mouse while outside of the interaction zone, was manually analyzed by an observer blind to treatment.

#### Inhibition of oxytocin synthesis within the BNST

To inhibit oxytocin synthesis, we used vivo-morpholinos (Genetools, LCC), which can prevent translation of a target sequence by blocking the translation initiation complex. The sequence used to block *Oxytocin* mRNA (antisense) was 5’-TTGGTGTTCTGAGTCCTCGATCC-3’, and 5’-TTCGTCTTCTGACTCCTCCATGC-3’ was used as a missense control. One week after the last episode of control handling or social defeat, animals were randomly assigned to receive one bilateral 0.2uL injection of 100pM morpholino missense or antisense aimed a little anterior to medioventral BNST (BNSTmv, AP:0.39, ML: 1.1, DV:-5.85). These coordinates were selected to minimize chances of spreading to the hypothalamic paraventricular nucleus (PVN). Animals were allowed to recover for one week before behavior testing. To confirm successful local inhibition of *Oxytocin* translation, 40 µm sections of the BNSTmv (3 slices per animal) and adjacent PVN (4 slices per animal) were stained with an oxytocin antibody (supplementary table 1). Nissl staining was used to assess tissue integrity (Reissner et al., 2012). For reagents and concentrations used, see supplementary table 1.

#### Oxytocin circuit visualization using recombinant adeno-associated virus (rAAV)

To identify projections of medioventral BNST oxytocin neurons, we used recombinant adeno-associated virus expressing Venus equipped with 2.6K mouse oxytocin promoter (AAV-OTpr-Venus) developed by the Grinevich lab (Knobloch et al., 2012). This construct has been successfully used for anatomical studies of oxytocin circuits in various rodent species including rats (Knobloch et al., 2012; Hasan et al., 2019; Tang et al.,in press), mice (Zheng et al., 2014), and prairie voles (Bosch et al., 2016). Males and females received a bilateral 40nl injection of AAV-OTpr-Venus (∼10^10^ copies per µl) into the BNSTmv. We chose to inject the virus anterior to the oxytocin neuron BNST population (AP:0.41, ML:1.1, DV:-5.85) to minimize chances of also infecting hypothalamic oxytocin neurons of the PVN. To assess successful anatomical targeting and specificity of infection, immunohistochemistry was used to detect Venus and oxytocin positive cells in 40um brain sections from the BNSTmv and PVN (see supplementary table 1 for antibodies). 1um z-stack images were then obtained using an Olympus FV1000 laser point-scanning confocal microscope.

#### Quantitative real-time PCR (qPCR) and fluorescent *in situ* hybridization

To assess the effects of social defeat stress on Oxtr expression in the anterior BNST, we used qPCR (supplementary table 3 for primers). Next, to assess whether the cells that express *Oxtr* in the BNSTam are excitatory or inhibitory, we performed fluorescent *in situ* hybridization (FISH) studies (ACDBio RNAscope^®^) using probes to detect *Oxtr, Gad1*, or *vGlut* based on California mouse mRNA sequences (for full methods and sequence information, see supplementary). We used an Olympus FV1000 laser point-scanning confocal microscope to collect 20x 1um z-stack images. Images where then analyzed for total expression and colocalization of *Oxtr*/*Gad1* or *Oxtr*/*vGlu2* using Fiji.

#### Effects of exogenous oxytocin infusion within anteromedial BNST (BNSTam)

To test whether oxytocin acting within the BNSTam is sufficient to induce social avoidance and vigilance, we injected one of two doses of synthetic oxytocin (Tocris) into the BNSTam of females and males naïve to stress. Mice were implanted with bilateral guide cannula (Plastics One, Roanoke, VA) using coordinates as previously published (Duque-Wilckens et al., 2018): AP: + 0.45, ML: +/−1.0, DV: + 4.6. Mice were single housed after surgery. After 7 days, mice were randomly assigned to receive bilateral 200nl infusions of either aCSF, 0.2ng (females only), or 1ng (males and females) of oxytocin using internal guides that projected 1 mm past the guides. Thirty minutes later, each mouse was tested in the social interaction test. Histology was used to determine injection sites as previously published (Duque-Wilckens et al., 2018).

### Experiments in C57BL/6 mice

#### Animals

All studies were approved by the Institutional Animal Care and Use Committee (IACUC) and conformed to NIH guidelines. Eight-week old female and male C57BL/6 mice (*Mus musculus*) and retired CD1 male breeder mice (*Mus musculus*) were purchased from Charles River (Hollister, CA). All mice were housed in standard polypropylene cages containing wood shavings. Experimental C57BL/6 mice were group housed (3-4 same sex animals), while CD1 mice, used as aggressors for vicarious defeat, were single housed. Mice were maintained in a colony room with a 12-h light/dark cycle (lights on at 7:00 h), 22°C, and with access to food (Harlan Teklad 7912; Madison, WI) and water *ad libitum*.

#### Vicarious defeat (c57BL/6 mice)

Female mice were randomly assigned to control handling or vicarious social defeat stress as described previously (Iñiguez et al., 2018). CD1 aggressors were housed in one side of a cage containing clear perforated acrylic glass dividers. Male C57BL/6 mice were placed for 10 minutes in the same side as the CD1 aggressor, where they were physically attacked. These males were not used for this study. Meanwhile, a female C57BL/6 mouse was placed in the neighboring side, experiencing the physical aggression vicariously through visual, olfactory, and auditory cues. Following each defeat episode, vicariously defeated animals were housed for 24h next to a novel CD1 aggressor until the next stress episode. This was repeated for 10 consecutive days. Control mice were housed in each side of a divided cage (two C57BL/6 mice per cage, same sex) and handled daily. Immediately after the last episode of vicarious defeat, all mice were single housed. Social interaction test was performed 24h later.

#### Social interaction test (C57BL/6 mice)

The social interaction test consisted of two phases lasting 150 seconds each (Iñiguez et al., 2018). In the first phase (target-absent), the animal was placed into an empty arena (40 cm × 40 cm × 40 cm) containing a circular wire cage on one side and was allowed to freely explore. In the second phase (target present), an unfamiliar male CD1 mouse was placed in the wire cage. Behavior was recorded via an automated video-tracking system (EthovisionXT; Noldus, Leesburg, VA). Vigilance behavior, defined as time spent with head oriented toward the target mouse while outside of the interaction zone (8-cm-wide corridor surrounding the wire cage), was manually analyzed by an observer blind to treatment.

#### Oxytocin immunohistochemistry in C57BL/6 mice exposed to vicarious defeat

To assess whether vicarious defeat induces changes in oxytocin activity within the BNSTmv, 40 µm brain sections of control and stressed females were stained for oxytocin (for antibodies used, see supplementary table 2).

#### Statistical analyses (all experiments)

All analyses were done using R software. 2-way ANOVAS were used to analyze effects of morpholino injections on female behavior and oxytocin neuron expression (treatment*stress), as well as the effects of vicarious defeat stress on C57BL/6 mice behavior (sex*stress). T-tests were used to assess acute effects of social defeat stress on male California mouse behavior and oxytocin+ cell detection in the BNSTmv of female C57BL/6. Male California mouse vigilance behavior was log transformed for analysis due to heterogeneous variance. Planned comparisons (package lsmeans in R [R Project for Statistical Computing, version 3.0.3, Vienna, Austria], Bonferroni, 0.95 confidence interval) were used if ANOVAS showed significant main or interaction effects. Effect size is reported as Cohen’s *d*.

## Acknowledgements

The authors thank C. J. Clayton for animal care and I. Brust-Mascher for imaging assistance. This work supported by Becas Chile Comisión Nacional de Investigación Científica y Tecnológica to ND-W, SC3GM130467 to SDI, DFG Collaborative Research Center (SFB) 1158, DFG grants GR 3619/701, GR 3619/4-1, SFB 1158, SNSF-DFG grant GR 3619/8-1, and Fritz Thyssen Foundation grant 10.16.2.018 MN to VG, and NIH R01 MH121829-01 to BCT. The authors report no biomedical financial interests or potential conflicts of interest.

## Supplementary Information

### Supplementary Methods

#### Quantitative Real-Time PCR

Anterior BNST was dissected from fresh frozen tissue and RNA was extracted using RNAeasy Mini Kits (Qiagen) and QIAzol as a lysis reagent (Qiagen). Reverse transcription was performed using iScript (BioRad) and SYBR green chemistry on an Applied Biosystems ViiA7 instrument was used to sequence amplification. Specific forward and reverse primers for *Oxtr* (Genbank accession: MN265350): and *B2m* (Genbank accession: XM_006995122) mRNA were designed based on California mouse sequence (see Supplementary Table 2). There were no differences in cycle thresholds between groups for *B2m*. Oxtr mRNA was normalized to B2m expression in each sample.

#### Fluorescent In-Situ Hybridization

Fluorescent in situ hybridization was performed following ACDBio RNAscope multiplex fluorescence methods. Probes were designed to detect California mouse Oxtr (2017-2961 of GFCW01069365.1), Gad1 (1188-2104 of GFCW01047509.1), and vGlut2 (675-1662 of GFCW01050557.1). Brains from adult male (n=2) and female (n=2) mice were flash frozen and cut at 20 μm. Sections were fixed in cold 10% neutral buffered formalin for 15 min and then dehydrated in a series of ethanol baths (50%, 70%, 100%), after which protease IV was applied to each section for 30 min. Sections where then washed in phosphate buffered saline (PBS). The probes were diluted 1:50 and applied to slides for 2 h at 40° C: one set of sections were probed for *Oxtr/Gad1* and the other set for *Oxtr/vGlu2*. Next, slides were rinsed in wash buffer and incubated sequentially in amplification buffer (AMP) 1 for 30 min at 40° C, then rinsed again in wash buffer and incubated in AMP 2 for 30 min at 40° C. Finally, slides were developed in Cy3 (*Oxtr*) for 15 min and then fluorescein (*Gad1 or vGlu2*) for 30 min, and coverslipped in Vectashield with DAPI. Z-stack images were collected with a 20x confocal. For colocalizations analyses, *Oxtr* nuclei were counted and then determined to be *Gad1* or *vGlu2* positive or negative.

**Supplementary Table 1.** Antibodies and reagents in California mouse tissue.

| Reagent | Dilution | Company (catalog #) |
| --- | --- | --- |
| Mouse anti-oxytocin antibody | 1:2000 | Sigma-Aldrich (MAB5296) |
| Chicken anti-GFP antibody | 1:1200 | Abcam (AB13970) |
| Goat anti-mouse biotinylated antibody | 1:250 | Vector (BA9200) |
| Goat anti-Chicken Alexafluor 488 antibody | 1:500 | Abcam (AB150169) |
| Streptavidin Alexa fluor 350 conjugate antibody | 1:500 | Thermofisher scientific (S11249) |
| **NeuroTrace™ 500/525 Green Fluorescent Nissl Stain** | 1:100 | Thermofisher scientific (N21480) |

**Supplementary Table 2.** Antibodies and reagents in C57/Balb6 tissue.

| Reagent | Dilution | Company (catalog #) |
| --- | --- | --- |
| Mouse anti-oxytocin antibody | 1:5000 | Sigma-Aldrich (MAB5296) |
| AffiniPure Fab Fragment Goat Anti-Mouse antibody | 1:25 | Jackson Laboratories (115-007-003) |
| Goat anti-mouse biotinylated antibody | 1:500 | Vector (BA9200) |
| Streptavidin Alexa fluor 555 conjugate antibody | 1:500 | Thermofisher scientific (S21381) |

**Supplementary Table 3.**
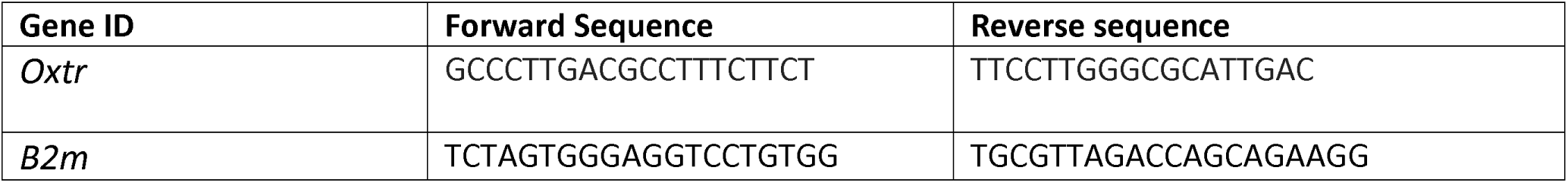
Sequences for qPCR primers.

**Supplementary figure 1.**
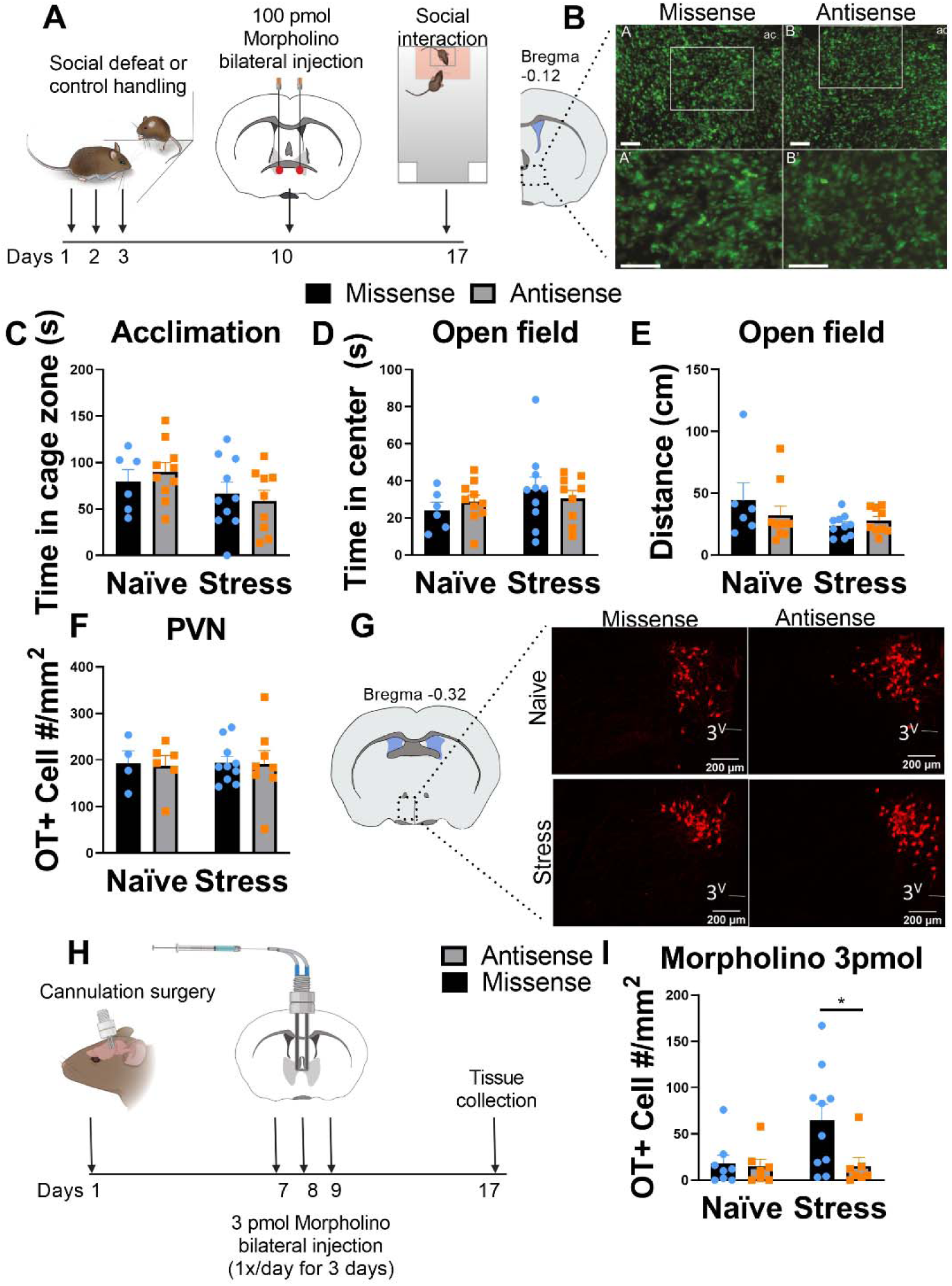
**A**. Timeline of experiment assessing effects of one bilateral 100pmol morpholino injections within BNSTmv on oxytocin expression and behavior. **B**. Representative image of Nissl stain used to assess tissue integrity after morpholino injections. **C**,**D**,**E**. There were no effects of antisense injections on approach, time spent in center, or distance traveled when no social target was present. **F**. Injections of antisense in the BNSTmv did not affect oxytocin cell numbers in the PVN. **G**. Representative photomicrographs of oxytocin+ cells in the PVN of naïve and stress females receiving morpholino missense and antisense. **H**. Timeline of experiment assessing effects of three daily bilateral injections of 3pmol morpholino injections within BNSTmv on oxytocin expression. **I**. Stress increased BNSTmv oxytocin cell number in animals receiving missense, but this was prevented by injections of morpholino missense.

**Supplementary figure 2.**
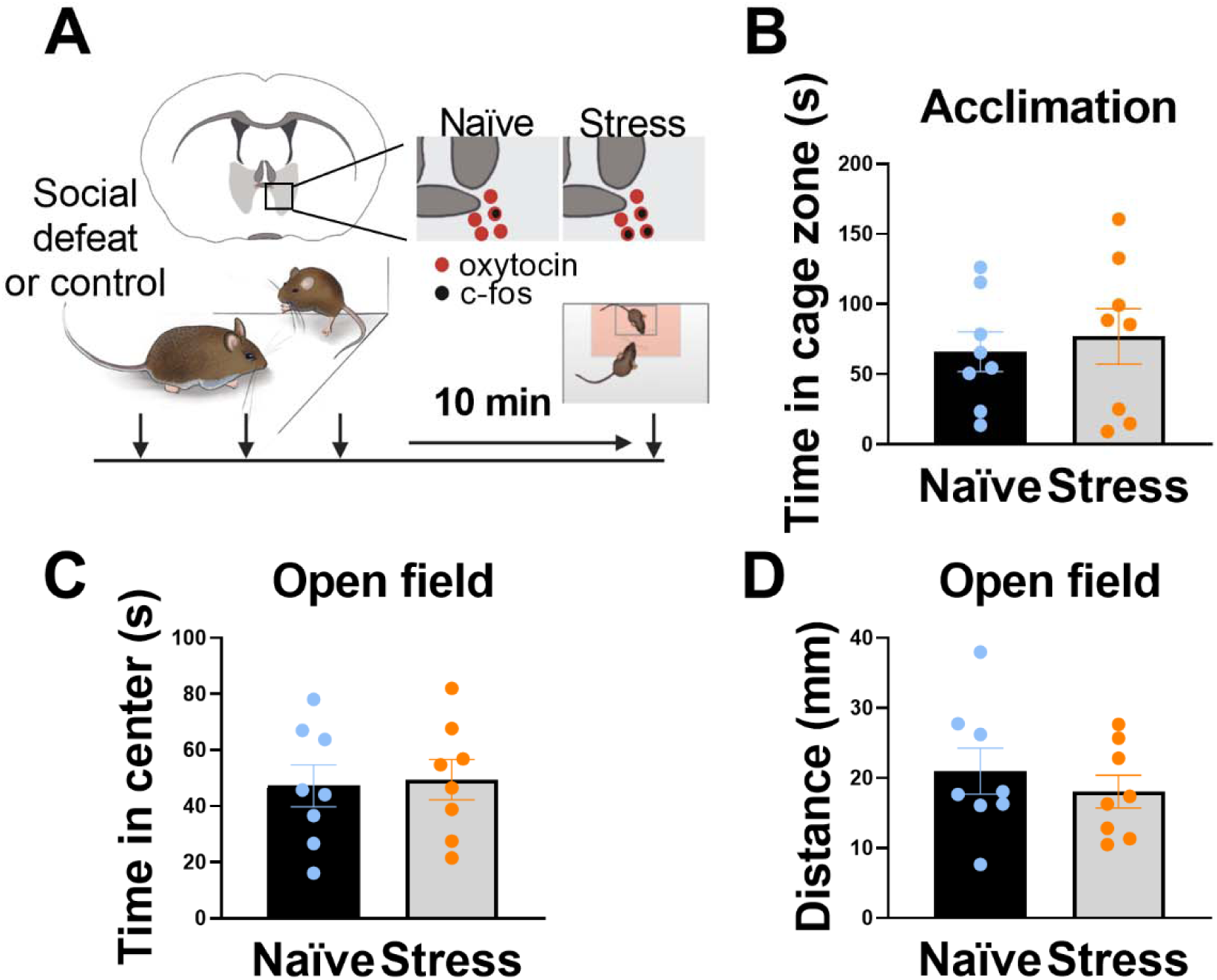
**A**. Timeline of experiment assessing the behavior of male California mice 10 min after a third episode of social defeat. **B**,**C**,**D**. Social defeat did not affect approach, time in center, or distance traveled when there was no social target present.

**Supplementary figure 3.**
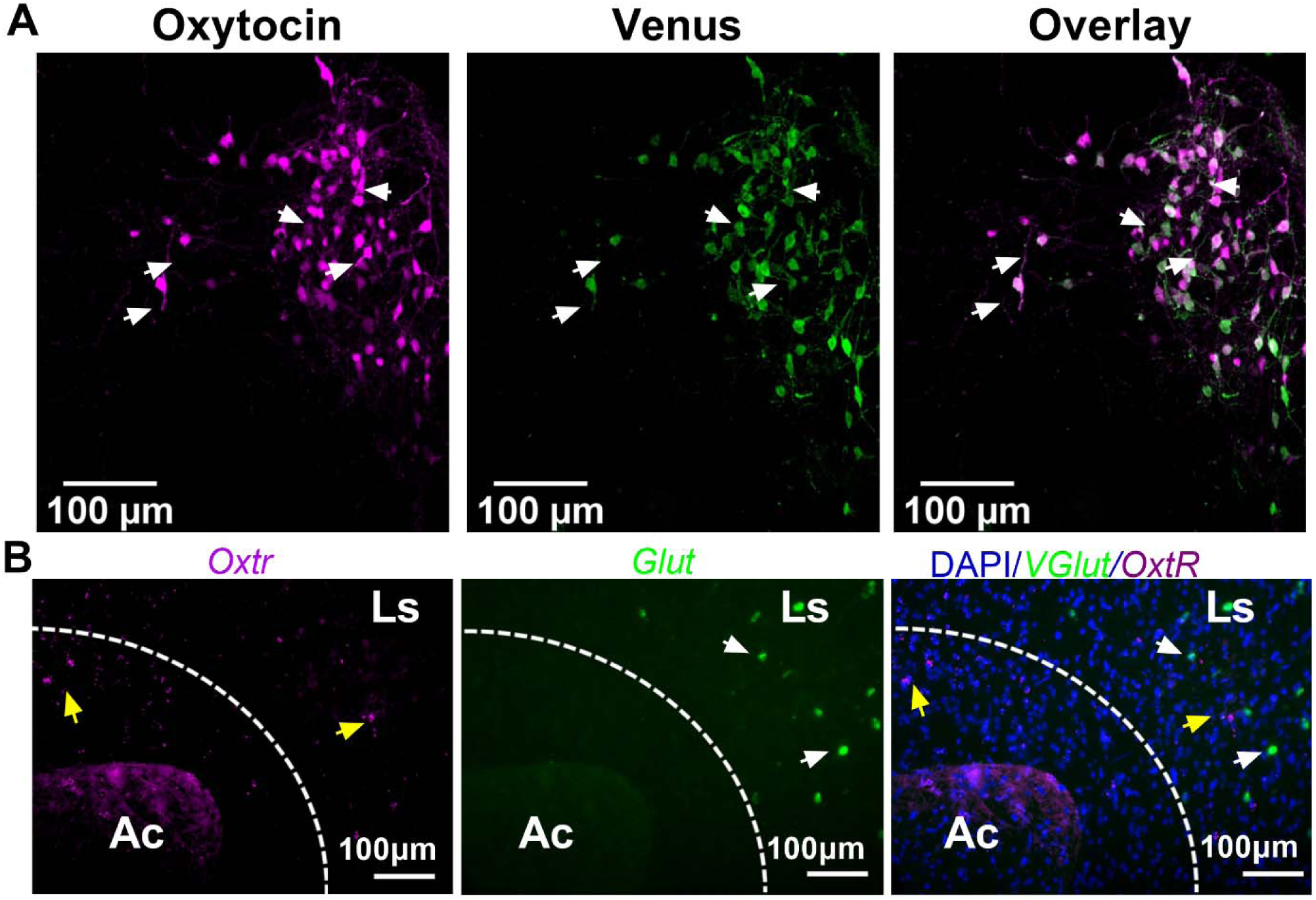
**A**. Representative photomicrographs of oxytocin, Venus, and colocalizations in the PVN after injection of OTpr AAV. **B**. Representative photomicrographs showing expression of OxtR and Glut. No Glut+ cells were found in the BNSTam, but there were Glut+ cells in the adjacent lateral septum. Ac=anterior commissure, Ls=lateral septum

## References

Allen, L.S., Gorski, R.A., 1990. Sex difference in the bed nucleus of the stria terminalis of the human brain. J. Comp. Neurol. 302, 697–706. https://doi.org/10.1002/cne.903020402

American Psychiatric Organization, 2013. Diagnostic and Statistical Manual of Mental Disorders (DSM-5^®^). American Psychiatric Pub.

Asher, M., Asnaani, A., Aderka, I.M., 2017. Gender differences in social anxiety disorder: A review. Clin. Psychol. Rev. 56, 1–12. https://doi.org/10.1016/j.cpr.2017.05.004

Avery, S.N., Clauss, J.A., Blackford, J.U., 2016. The Human BNST: Functional Role in Anxiety and Addiction. Neuropsychopharmacology 41, 126–141. https://doi.org/10.1038/npp.2015.185

Bartz, J.A., Zaki, J., Bolger, N., Ochsner, K.N., 2011. Social effects of oxytocin in humans: context and person matter. Trends Cogn. Sci. 15, 301–309. https://doi.org/10.1016/j.tics.2011.05.002

Beesdo-Baum, K., Knappe, S., Fehm, L., Höfler, M., Lieb, R., Hofmann, S.G., Wittchen, H.-U., 2012. The natural course of social anxiety disorder among adolescents and young adults. Acta Psychiatr. Scand. 126, 411–425. https://doi.org/10.1111/j.1600-0447.2012.01886.x

Blume, A., Bosch, O.J., Miklos, S., Torner, L., Wales, L., Waldherr, M., Neumann, I.D., 2008. Oxytocin reduces anxiety via ERK1/2 activation: local effect within the rat hypothalamic paraventricular nucleus. Eur. J. Neurosci. 27, 1947–1956. https://doi.org/10.1111/j.1460-9568.2008.06184.x

Bosch, O.J., Dabrowska, J., Modi, M.E., Johnson, Z.V., Keebaugh, A.C., Barrett, C.E., Ahern, T.H., Guo, J., Grinevich, V., Rainnie, D.G., Neumann, I.D., Young, L.J., 2016. Oxytocin in the nucleus accumbens shell reverses CRFR2-evoked passive stress-coping after partner loss in monogamous male prairie voles. Psychoneuroendocrinology 64, 66–78. https://doi.org/10.1016/j.psyneuen.2015.11.011

Bosch, O.J., Young, L.J., 2018. Oxytocin and Social Relationships: From Attachment to Bond Disruption. Curr. Top. Behav. Neurosci. 35, 97–117. https://doi.org/10.1007/7854_2017_10

Caffé, A.R., Ryen, P.C.V., Woude, T.P.V.D., Leeuwen, F.W.V., 1989. Vasopressin and oxytocin systems in the brain and upper spinal cord of Macaca fascicularis. J. Comp. Neurol. 287, 302–325. https://doi.org/10.1002/cne.902870304

Campi, K.L., Jameson, C.E., Trainor, B.C., 2013. Sexual Dimorphism in the Brain of the Monogamous California Mouse (Peromyscus californicus). Brain. Behav. Evol. 81, 236–249. https://doi.org/10.1159/000353260

Chen, N.T.M., Clarke, P.J.F., 2017. Gaze-Based Assessments of Vigilance and Avoidance in Social Anxiety: a Review. Curr. Psychiatry Rep. 19, 59. https://doi.org/10.1007/s11920-017-0808-4

Chini, B., Verhage, M., Grinevich, V., 2017. The Action Radius of Oxytocin Release in the Mammalian CNS: From Single Vesicles to Behavior. Trends Pharmacol. Sci. 38, 982–991. https://doi.org/10.1016/j.tips.2017.08.005

Clauss, J.A., Avery, S.N., Benningfield, M.M., Blackford, J.U., 2019. Social anxiety is associated with BNST response to unpredictability. Depress. Anxiety 36, 666–675. https://doi.org/10.1002/da.22891

Delville, Y., De Vries, G.J., Ferris, C.F., 2000. Neural connections of the anterior hypothalamus and agonistic behavior in golden hamsters. Brain. Behav. Evol. 55, 53–76. https://doi.org/10.1159/000006642

DiBenedictis, B.T., Nussbaum, E.R., Cheung, H.K., Veenema, A.H., 2017. Quantitative mapping reveals age and sex differences in vasopressin, but not oxytocin, immunoreactivity in the rat social behavior neural network. J. Comp. Neurol. https://doi.org/10.1002/cne.24216

Dölen, G., Darvishzadeh, A., Huang, K.W., Malenka, R.C., 2013. Social reward requires coordinated activity of nucleus accumbens oxytocin and serotonin. Nature 501, 179–184. https://doi.org/10.1038/nature12518

Domes, G., Heinrichs, M., Gläscher, J., Büchel, C., Braus, D.F., Herpertz, S.C., 2007. Oxytocin attenuates amygdala responses to emotional faces regardless of valence. Biol. Psychiatry 62, 1187–1190. https://doi.org/10.1016/j.biopsych.2007.03.025

Dong, H.-W., Swanson, L.W., 2004. Organization of axonal projections from the anterolateral area of the bed nuclei of the stria terminalis. J. Comp. Neurol. 468, 277–298. https://doi.org/10.1002/cne.10949

Duque-Wilckens, N., Steinman, M.Q., Busnelli, M., Chini, B., Yokoyama, S., Pham, M., Laredo, S.A., Hao, R., Perkeybile, A.M., Minie, V.A., Tan, P.B., Bales, K.L., Trainor, B.C., 2018. Oxytocin Receptors in the Anteromedial Bed Nucleus of the Stria Terminalis Promote Stress-Induced Social Avoidance in Female California Mice. Biol. Psychiatry 83, 203–213. https://doi.org/10.1016/j.biopsych.2017.08.024

Duque-Wilckens, N., Steinman, M.Q., Laredo, S.A., Hao, R., Perkeybile, A.M., Bales, K.L., Trainor, B.C., 2016. Inhibition of vasopressin V1a receptors in the medioventral bed nucleus of the stria terminalis has sex- and context-specific anxiogenic effects. Neuropharmacology 110, 59–68. https://doi.org/10.1016/j.neuropharm.2016.07.018

Eckstein, M., Scheele, D., Weber, K., Stoffel-Wagner, B., Maier, W., Hurlemann, R., 2014. Oxytocin facilitates the sensation of social stress. Hum. Brain Mapp. 35, 4741–4750. https://doi.org/10.1002/hbm.22508

Figel, B., Brinkmann, L., Buff, C., Heitmann, C.Y., Hofmann, D., Bruchmann, M., Becker, M.P.I., Herrmann, M.J., Straube, T., 2019. Phasic amygdala and BNST activation during the anticipation of temporally unpredictable social observation in social anxiety disorder patients. NeuroImage Clin. 22, 101735. https://doi.org/10.1016/j.nicl.2019.101735

Fox, A.S., Shackman, A.J., 2019. The central extended amygdala in fear and anxiety: Closing the gap between mechanistic and neuroimaging research. Neurosci. Lett. 693, 58–67. https://doi.org/10.1016/j.neulet.2017.11.056

Goode, T.D., Maren, S., 2017. Role of the bed nucleus of the stria terminalis in aversive learning and memory. Learn. Mem. Cold Spring Harb. N 24, 480–491. https://doi.org/10.1101/lm.044206.116

Greenberg, G.D., Laman-Maharg, A., Campi, K.L., Voigt, H., Orr, V.N., Schaal, L., Trainor, B.C., 2014. Sex differences in stress-induced social withdrawal: role of brain derived neurotrophic factor in the bed nucleus of the stria terminalis. Front. Behav. Neurosci. 7, 223. https://doi.org/10.3389/fnbeh.2013.00223

Greenberg, G.D., Laman-Maharg, A., Campi, K.L., Voigt, H., Orr, V.N., Schaal, L., Trainor, B.C., 2013. Sex differences in stress-induced social withdrawal: role of brain derived neurotrophic factor in the bed nucleus of the stria terminalis. Front. Behav. Neurosci. 7, 223. https://doi.org/10.3389/fnbeh.2013.00223

Grinevich, V., Desarménien, M.G., Chini, B., Tauber, M., Muscatelli, F., 2015. Ontogenesis of oxytocin pathways in the mammalian brain: late maturation and psychosocial disorders. Front. Neuroanat. 8. https://doi.org/10.3389/fnana.2014.00164

Grinevich, V., Knobloch-Bollmann, H.S., Eliava, M., Busnelli, M., Chini, B., 2016. Assembling the Puzzle: Pathways of Oxytocin Signaling in the Brain. Biol. Psychiatry, Oxytocin and Psychiatry: From DNA to Social Behavior 79, 155–164. https://doi.org/10.1016/j.biopsych.2015.04.013

Han, R.T., Kim, Y.-B., Park, E.-H., Kim, J.Y., Ryu, C., Kim, H.Y., Lee, J., Pahk, K., Shanyu, C., Kim, H., Back, S.K., Kim, H.J., Kim, Y.I., Na, H.S., 2018. Long-Term Isolation Elicits Depression and Anxiety-Related Behaviors by Reducing Oxytocin-Induced GABAergic Transmission in Central Amygdala. Front. Mol. Neurosci. 11, 246. https://doi.org/10.3389/fnmol.2018.00246

Hatton, G.I., 1990. Emerging concepts of structure-function dynamics in adult brain: the hypothalamo-neurohypophysial system. Prog. Neurobiol. 34, 437–504. https://doi.org/10.1016/0301-0082(90)90017-b

Heimberg, R.G., 1995. Social Phobia: Diagnosis, Assessment, and Treatment. Guilford Press.

Hodes, G.E., Pfau, M.L., Purushothaman, I., Ahn, H.F., Golden, S.A., Christoffel, D.J., Magida, J., Brancato, A., Takahashi, A., Flanigan, M.E., Menard, C., Aleyasin, H., Koo, J.W., Lorsch, Z.S., Feng, J., Heshmati, M., Wang, M., Turecki, G., Neve, R., Zhang, B., Shen, L., Nestler, E.J., Russo, S.J., 2015. Sex differences in nucleus accumbens transcriptome profiles associated with susceptibility versus resilience to subchronic variable stress. J Neurosci 35, 16362–16376.

Hung, L.W., Neuner, S., Polepalli, J.S., Beier, K.T., Wright, M., Walsh, J.J., Lewis, E.M., Luo, L., Deisseroth, K., Dölen, G., Malenka, R.C., 2017. Gating of social reward by oxytocin in the ventral tegmental area. Science 357, 1406–1411. https://doi.org/10.1126/science.aan4994

Iñiguez, S.D., Flores-Ramirez, F.J., Riggs, L.M., Alipio, J.B., Garcia-Carachure, I., Hernandez, M.A., Sanchez, D.O., Lobo, M.K., Serrano, P.A., Braren, S.H., Castillo, S.A., 2018. Vicarious Social Defeat Stress Induces Depression-Related Outcomes in Female Mice. Biol. Psychiatry, Novel Mechanisms of Antidepressant Action 83, 9–17. https://doi.org/10.1016/j.biopsych.2017.07.014

Ipser, J.C., Kariuki, C.M., Stein, D.J., 2008. Pharmacotherapy for social anxiety disorder: a systematic review. Expert Rev. Neurother. 8, 235–257. https://doi.org/10.1586/14737175.8.2.235

Janitzky, K., Peine, A., Kröber, A., Yanagawa, Y., Schwegler, H., Roskoden, T., 2014. Increased CRF mRNA expression in the sexually dimorphic BNST of male but not female GAD67 mice and TMT predator odor stress effects upon spatial memory retrieval. Behav. Brain Res. 272, 141–149. https://doi.org/10.1016/j.bbr.2014.06.020

Kessler, R.C., Berglund, P., Demler, O., Jin, R., Merikangas, K.R., Walters, E.E., 2005. Lifetime prevalence and age-of-onset distributions of DSM-IV disorders in the National Comorbidity Survey Replication. Arch. Gen. Psychiatry 62, 593–602. https://doi.org/10.1001/archpsyc.62.6.593

Kirsch, P., Esslinger, C., Chen, Q., Mier, D., Lis, S., Siddhanti, S., Gruppe, H., Mattay, V.S., Gallhofer, B., Meyer-Lindenberg, A., 2005. Oxytocin Modulates Neural Circuitry for Social Cognition and Fear in Humans. J. Neurosci. 25, 11489–11493. https://doi.org/10.1523/JNEUROSCI.3984-05.2005

Knobloch, H.S., Charlet, A., Hoffmann, L.C., Eliava, M., Khrulev, S., Cetin, A.H., Osten, P., Schwarz, M.K., Seeburg, P.H., Stoop, R., Grinevich, V., 2012. Evoked Axonal Oxytocin Release in the Central Amygdala Attenuates Fear Response. Neuron 73, 553–566. https://doi.org/10.1016/j.neuron.2011.11.030

Knobloch, H.S., Grinevich, V., 2014. Evolution of oxytocin pathways in the brain of vertebrates. Front. Behav. Neurosci. 8. https://doi.org/10.3389/fnbeh.2014.00031

Kollack-Walker, S., Watson, S.J., Akil, H., 1997. Social Stress in Hamsters: Defeat Activates Specific Neurocircuits within the Brain. J. Neurosci. 17, 8842–8855. https://doi.org/10.1523/JNEUROSCI.17-22-08842.1997

Landgraf, R., Neumann, I.D., 2004. Vasopressin and oxytocin release within the brain: a dynamic concept of multiple and variable modes of neuropeptide communication. Front. Neuroendocrinol. 25, 150–176. https://doi.org/10.1016/j.yfrne.2004.05.001

Lebow, M.A., Chen, A., 2016. Overshadowed by the amygdala: the bed nucleus of the stria terminalis emerges as key to psychiatric disorders. Mol. Psychiatry 21, 450–463. https://doi.org/10.1038/mp.2016.1

Liebowitz, M.R., Gelenberg, A.J., Munjack, D., 2005. Venlafaxine extended release vs placebo and paroxetine in social anxiety disorder. Arch. Gen. Psychiatry 62, 190–198. https://doi.org/10.1001/archpsyc.62.2.190

Markham, C.M., Norvelle, A., Huhman, K.L., 2009. Role of the bed nucleus of the stria terminalis in the acquisition and expression of conditioned defeat in Syrian hamsters. Behav. Brain Res. 198, 69–73. https://doi.org/10.1016/j.bbr.2008.10.022

Marlin, B.J., Mitre, M., D’amour, J.A., Chao, M.V., Froemke, R.C., 2015. Oxytocin enables maternal behaviour by balancing cortical inhibition. Nature 520, 499–504. https://doi.org/10.1038/nature14402

Martinon, D., Lis, P., Roman, A.N., Tornesi, P., Applebey, S.V., Buechner, G., Olivera, V., Dabrowska, J., 2019. Oxytocin receptors in the dorsolateral bed nucleus of the stria terminalis (BNST) bias fear learning toward temporally predictable cued fear. Transl. Psychiatry 9, 140. https://doi.org/10.1038/s41398-019-0474-x

Maynard, K.R., Hobbs, J.W., Phan, B.N., Gupta, A., Rajpurohit, S., Williams, C., Rajpurohit, A., Shin, J.H., Jaffe, A.E., Martinowich, K., 2018. BDNF-TrkB signaling in oxytocin neurons contributes to maternal behavior. eLife 7. https://doi.org/10.7554/eLife.33676

Mittaud, P., Labourdette, G., Zingg, H., Scala, D.G.-D., 2002. Neurons modulate oxytocin receptor expression in rat cultured astrocytes: Involvement of TGF-β and membrane components. Glia 37, 169–177. https://doi.org/10.1002/glia.10029

Moaddab, M., Dabrowska, J., 2017. Oxytocin receptor neurotransmission in the dorsolateral bed nucleus of the stria terminalis facilitates the acquisition of cued fear in the fear-potentiated startle paradigm in rats. Neuropharmacology 121, 130–139. https://doi.org/10.1016/j.neuropharm.2017.04.039

Nasanbuyan, N., Yoshida, M., Takayanagi, Y., Inutsuka, A., Nishimori, K., Yamanaka, A., Onaka, T., 2018. Oxytocin-Oxytocin Receptor Systems Facilitate Social Defeat Posture in Male Mice. Endocrinology 159, 763–775. https://doi.org/10.1210/en.2017-00606

Newman, E.L., Covington, H.E., Suh, J., Bicakci, M.B., Ressler, K.J., DeBold, J.F., Miczek, K.A., 2019. Fighting Females: Neural and Behavioral Consequences of Social Defeat Stress in Female Mice. Biol. Psychiatry 86, 657–668. https://doi.org/10.1016/j.biopsych.2019.05.005

Park, H., Poo, M., 2013. Neurotrophin regulation of neural circuit development and function. Nat. Rev. Neurosci. 14, 7–23. https://doi.org/10.1038/nrn3379

Reissner, K.J., Sartor, G.C., Vazey, E.M., Dunn, T.E., Aston-Jones, G., Kalivas, P.W., 2012. Use of vivo-morpholinos for control of protein expression in the adult rat brain. J. Neurosci. Methods 203, 354–360. https://doi.org/10.1016/j.jneumeth.2011.10.009

Rogers-Carter, M.M., Varela, J.A., Gribbons, K.B., Pierce, R.C., McGoey, M.T., Ritchey, M., Christianson, J.P., 2018. Insular cortex mediates approach and avoidance responses to social affective stimuli. Nat. Neurosci. in press.

Ross, H.E., Cole, C.D., Smith, Y., Neumann, I.D., Landgraf, R., Murphy, A.Z., Young, L.J., 2009. Characterization of the oxytocin system regulating affiliative behavior in female prairie voles. Neuroscience 162, 892–903. https://doi.org/10.1016/j.neuroscience.2009.05.055

Ruscio, A.M., Brown, T.A., Chiu, W.T., Sareen, J., Stein, M.B., Kessler, R.C., 2008. Social fears and social phobia in the USA: results from the National Comorbidity Survey Replication. Psychol. Med. 38, 15–28. https://doi.org/10.1017/S0033291707001699

Shamay-Tsoory, S.G., Abu-Akel, A., 2016. The Social Salience Hypothesis of Oxytocin. Biol. Psychiatry, Oxytocin and Psychiatry: From DNA to Social Behavior 79, 194–202. https://doi.org/10.1016/j.biopsych.2015.07.020

Song, Z., Borland, J.M., Larkin, T.E., O’Malley, M., Albers, H.E., 2016. Activation of oxytocin receptors, but not arginine-vasopressin V1a receptors, in the ventral tegmental area of male Syrian hamsters is essential for the reward-like properties of social interactions. Psychoneuroendocrinology 74, 164–172. https://doi.org/10.1016/j.psyneuen.2016.09.001

Spence, S.H., Rapee, R.M., 2016. The etiology of social anxiety disorder: An evidence-based model. Behav. Res. Ther. 86, 50–67. https://doi.org/10.1016/j.brat.2016.06.007

Stein, D.J., Lim, C.C.W., Roest, A.M., de Jonge, P., Aguilar-Gaxiola, S., Al-Hamzawi, A., Alonso, J., Benjet, C., Bromet, E.J., Bruffaerts, R., de Girolamo, G., Florescu, S., Gureje, O., Haro, J.M., Harris, M.G., He, Y., Hinkov, H., Horiguchi, I., Hu, C., Karam, A., Karam, E.G., Lee, S., Lepine, J.-P., Navarro-Mateu, F., Pennell, B.-E., Piazza, M., Posada-Villa, J., Ten Have, M., Torres, Y., Viana, M.C., Wojtyniak, B., Xavier, M., Kessler, R.C., Scott, K.M., WHO World Mental Health Survey Collaborators, 2017. The cross-national epidemiology of social anxiety disorder: Data from the World Mental Health Survey Initiative. BMC Med. 15, 143. https://doi.org/10.1186/s12916-017-0889-2

Steinman, M.Q., Duque-Wilckens, N., Greenberg, G.D., Hao, R., Campi, K.L., Laredo, S.A., Laman-Maharg, A., Manning, C.E., Doig, I.E., Lopez, E.M., Walch, K., Bales, K.L., Trainor, B.C., 2016. Sex-Specific Effects of Stress on Oxytocin Neurons Correspond With Responses to Intranasal Oxytocin. Biol. Psychiatry, Corticotropin-Releasing Factor, FKBP5, and Posttraumatic Stress Disorder 80, 406–414. https://doi.org/10.1016/j.biopsych.2015.10.007

Steinman, M.Q., Duque-Wilckens, N., Trainor, B.C., 2019. Complementary Neural Circuits for Divergent Effects of Oxytocin: Social Approach Versus Social Anxiety. Biol. Psychiatry, Longitudinal Perspectives on Stress and Depression 85, 792–801. https://doi.org/10.1016/j.biopsych.2018.10.008

Stern, J.E., Armstrong, W.E., 1998. Reorganization of the dendritic trees of oxytocin and vasopressin neurons of the rat supraoptic nucleus during lactation. J. Neurosci. Off. J. Soc. Neurosci. 18, 841–853.

Takahashi, A., Chung, J.-R., Zhang, S., Zhang, H., Grossman, Y., Aleyasin, H., Flanigan, M.E., Pfau, M.L., Menard, C., Dumitriu, D., Hodes, G.E., McEwen, B.S., Nestler, E.J., Han, M.-H., Russo, S.J., 2017. Establishment of a repeated social defeat stress model in female mice. Sci. Rep. 7, 12838. https://doi.org/10.1038/s41598-017-12811-8

Tang, Y., Benusiglio, D., Lefevre, A., Hilfiger, L., Althammer, F., Bludau, A., Hagiwara, D., Baudon, A., Darbon, P., Shimmer, J., Kirchner, M.K., Roy, R.K., Wang, S., Eliava, M., Wagner, S., Oberhuber, M., Conzelmann, K.K., Schwarz, M., Stern, J.E., Leng, G., Neumann, I.D., Charlet, A., Grinevich, V., 2020. Social touch promotes inter-female communication via activation of oxytocin parvocellular neurons. Nature Neurosci.

Theodosis, D.T., Piet, R., Poulain, D.A., Oliet, S.H.R., 2004. Neuronal, glial and synaptic remodeling in the adult hypothalamus: functional consequences and role of cell surface and extracellular matrix adhesion molecules. Neurochem. Int. 45, 491–501. https://doi.org/10.1016/j.neuint.2003.11.003

Trainor, B.C., Pride, M.C., Villalon Landeros, R., Knoblauch, N.W., Takahashi, E.Y., Silva, A.L., Crean, K.K., 2011. Sex differences in social interaction behavior following social defeat stress in the monogamous California mouse (Peromyscus californicus). PloS One 6, e17405. https://doi.org/10.1371/journal.pone.0017405

Trainor, B.C., Takahashi, E.Y., Campi, K.L., Florez, S.A., Greenberg, G.D., Laman-Maharg, A., Laredo, S.A., Orr, V.N., Silva, A.L., Steinman, M.Q., 2013. Sex differences in stress-induced social withdrawal: independence from adult gonadal hormones and inhibition of female phenotype by corncob bedding. Horm. Behav. 63, 543–550. https://doi.org/10.1016/j.yhbeh.2013.01.011

Van Ameringen, M.A., Lane, R.M., Walker, J.R., Bowen, R.C., Chokka, P.R., Goldner, E.M., Johnston, D.G., Lavallee, Y.J., Nandy, S., Pecknold, J.C., Hadrava, V., Swinson, R.P., 2001. Sertraline treatment of generalized social phobia: a 20-week, double-blind, placebo-controlled study. Am. J. Psychiatry 158, 275–281. https://doi.org/10.1176/appi.ajp.158.2.275

Veenema, A.H., Neumann, I.D., 2008. Central vasopressin and oxytocin release: regulation of complex social behaviours. Prog. Brain Res. 170, 261–276. https://doi.org/10.1016/S0079-6123(08)00422-6

Wang, Z., Aragona, B.J., 2004. Neurochemical regulation of pair bonding in male prairie voles. Physiol. Behav., Male Sexual Function 83, 319–328. https://doi.org/10.1016/j.physbeh.2004.08.024

Warren, B.L., Mazei-Robison, M., Robison, A.J., Iñiguez, S.D., 2020. Can I get a witness? Using vicarious defeat stress to study mood-related illnesses in traditionally understudied populations. Biol. Psychiatry 0. https://doi.org/10.1016/j.biopsych.2020.02.004

Welch, J.D., Kozareva, V., Ferreira, A., Vanderburg, C., Martin, C., Macosko, E.Z., 2019. Single-Cell Multiomic Integration Compares and Contrasts Features of Brain Cell Identity. Cell 177, 1873-1887.e17. https://doi.org/10.1016/j.cell.2019.05.006

Whylings, J., Rigney, N., Peters, N.V., de Vries, G.J., Petrulis, A., 2020. Sexually dimorphic role of BNST vasopressin cells in sickness and social behavior in male and female mice. Brain. Behav. Immun. 83, 68–77. https://doi.org/10.1016/j.bbi.2019.09.015

Williams, A.V., Duque-Wilckens, N., Ramos-Maciel, S., Campi, K.L., Bhela, S.K., Xu, C.K., Jackson, K., Chini, B., Pesavento, P.A., Trainor, B.C., 2020. Social approach and social vigilance are differentially regulated by oxytocin receptors in the nucleus accumbens. Neuropsychopharmacology 1–8. https://doi.org/10.1038/s41386-020-0657-4

Williams, A.V., Laman-Maharg, A., Armstrong, C.V., Ramos-Maciel, S., Minie, V.A., Trainor, B.C., 2018. Acute inhibition of kappa opioid receptors before stress blocks depression-like behaviors in California mice. Prog. Neuropsychopharmacol. Biol. Psychiatry 86, 166–174. https://doi.org/10.1016/j.pnpbp.2018.06.001

Zheng, J.-J., Li, S.-J., Zhang, X.-D., Miao, W.-Y., Zhang, D., Yao, H., Yu, X., 2014. Oxytocin mediates early experience–dependent cross-modal plasticity in the sensory cortices. Nat. Neurosci. 17, 391–399. https://doi.org/10.1038/nn.3634

